# Detection and characterization of copy number variants based on whole-genome sequencing by DNBSEQ platforms

**DOI:** 10.1101/786962

**Authors:** Junhua Rao, Lihua Peng, Fang Chen, Hui Jiang, Chunyu Geng, Xia Zhao, Xin Liu, Xinming Liang, Feng Mu

**Affiliations:** MGI, BGI-Shenzhen, Shenzhen 518083, China; BGI-Shenzhen, Shenzhen 518083, China; BGI-Qingdao, BGI-Shenzhen, Qingdao 266555 Shandong, China; MGI-Wuhan, BGI-Shenzhen, Wuhan 430074, China

## Abstract

**Background:** Next-generation sequence (NGS) has rapidly developed in past years which makes whole-genome sequencing (WGS) becoming a more cost- and time-efficient choice in wide range of biological researches. We usually focus on some variant detection via WGS data, such as detection of single nucleotide polymorphism (SNP), insertion and deletion (Indel) and copy number variant (CNV), which playing an important role in many human diseases. However, the feasibility of CNV detection based on WGS by DNBSEQ™ platforms was unclear. We systematically analysed the genome-wide CNV detection power of DNBSEQ™ platforms and Illumina platforms on NA12878 with five commonly used tools, respectively.

**Results:** DNBSEQ™ platforms showed stable ability to detect slighter more CNVs on genome-wide (average 1.24-fold than Illumina platforms). Then, CNVs based on DNBSEQ™ platforms and Illumina platforms were evaluated with two public benchmarks of NA12878, respectively. DNBSEQ™ and Illumina platforms showed similar sensitivities and precisions on both two benchmarks. Further, the difference between tools for CNV detection was analyzed, and indicated the selection of tool for CNV detection could affected the CNV performance, such as count, distribution, sensitivity and precision.

**Conclusion:** The major contribution of this paper is providing a comprehensive guide for CNV detection based on WGS by DNBSEQ™ platforms for the first time.

## Background

Copy number variation (CNV) is becoming more and more important in genomic variation since first reported in 2004[1, 2]. Currently, the large structure variation, especially CNV, have been heavily studied to clearly demonstrated them playing an important role in human diseases, such as autism[3-6], schizophrenia[7], Parkinson[8], Hirschsprung[9] and cancer[10]. In the same time, many sequencing technologies and methods about CNV come into being. Fluorescence *in situ* hybridization (FISH), array-based comparative genomic hybridization (array CGH)[11, 12] and SNP array were used for CNV detection at the beginning. Later, whole-exome sequencing (WES) and whole-genome sequencing (WGS), two methods based on next-generation sequencing (NGS), were introduced for CNV detection and were widely used in a large number of biological researches. Without the limitations of specified target regions associated with hybridization or array, WES and WGS become more advisable in CNV detection.

Illumina developed many sequencing platforms using reversible terminator chemistry and Illumina’s sequencing by synthesis (SBS) technology, such as HiSeq2500, HiSeq X Ten and NovaSeq6000[13]. WES and WGS sequenced on Illumina platforms were widely used in biological research, including CNV detection. Meanwhile, many tools were developed for CNV detection with WES[14] or WGS[15, 16]. There are five main strategies used in tools for CNV detection: (1) read pair (RP), (2) read depth (RD), (3) split read (SR), (4) de novo assembly (AS), and (5) combination of approaches (CA). And each strategy has its own advantage and limitation. RD-based tools can get accurate copy number but have no ability to detect the breakpoints of CNV. SR-based tools can get breakpoints at base-pair resolution but have low ability in low-complexity regions. AS-based tools could detect CNVs without a known reference but have high-requirement of computational resources [15]. Moreover, different tools were designed for different sample conditions. For example, BreakDancer[17] was developed for RP strategy and was built for single sample, HYDRA[18] based on CA strategy was suit for multiple samples and CNVnorm[19] based on RD strategy was designed for case-control condition.

MGI Tech Co., Ltd. (MGI), a subsidiary of BGI Group and the parent company of Complete Genomics™(CG), which is the instruments manufacturing business of BGI, had launched several platforms using DNBSEQ™ sequencing technology since 2016, such as BGISEQ-500, DNBSEQ-G400 (MGISEQ-2000) and DNBSEQ-T7. Compared to other existing sequencing platform, DNBSEQ™ sequencing technology combines the advantages of low amplification error rates from DNA nanoballs (DNBs), high density patterned array and combinational probe anchor synthesis (cPAS) sequencing technology. These advantages significantly improve sequencing accuracy and have much lower duplication rates and reduced index hopping. BGISEQ-500 platform showed high performance on detection of SNP and Indel on WGS[20, 21] or WES[22]. Furthermore, DNBSEQ™ platform is a good choice for transcriptome analysis in plants[23].

Accompanied by the improvement of NGS, primarily for decrease of time and cost, increase of data size per run and sequence quality, more CNV researches are carried out based on WGS by NGS platforms. Many CNV detection analysis based on WGS by Illumina platforms were evaluated, but there is no comprehensive guide for CNV detection based on WGS by DNBSEQ™ platform. Here, we presented a performance analysis of CNV detection by analysing eight WGS demo data sequenced on two DNBSEQ™ platforms with five common tools based on four strategies. For comparison, two Illumina platforms were used as control platforms and we indicated that both the sensitivity and the precision of CNVs based on DNBSEQ™ platforms were consistent with those of CNVs based on Illumina platforms.

## Results

### Selection of tools and benchmarks

We selected five tools for CNV detection from published CNV tools in three factors: single sample pattern, widely used and keep date (Supplementary Table S1). These five tools were designed for genome-wide CNV detection based on different strategies: BreakDancer[17] was built on RP, CNVnator[24] was built on RD, Pindel[25] was built on SR, DELLY[26] and LUMPY[27] were built on CA.

Moreover, two available CNV benchmarks of NA12878 were introduced for further evaluation of CNV: 2,819 CNVs from Layer, et al. in 2014 (LUMPY_2014)[27] and 2,171 CNVs from the 1000 Genome Project WGS data in 2015 (1KG_2015)[28]. A CNV in benchmark would be considered as specific if the proportion of its region size which overlapped with other CNVs in benchmark reciprocally was less than a threshold. When the threshold was 90.00%, 56.05% of CNVs in LUMPY_2014 and 32.43% of CNVs in 1KG_2015 were specific, respectively (Supplementary Figure S1a). While the threshold was 50%, the ratio of specific CNVs was 47.82% in LUMPY_2014 and 30.49% in 1KG_2015, respectively (Supplementary Figure S1b). In a word, there were large proportion specific CNVs in both LUMPY_2014 and 1KG_2015, so these two benchmarks both were used in this study.

### CNV detection and filtration

To detect CNV, we used ten WGS datasets of NA12878 sequenced on two DNBSEQ™ sequencing platforms (three datasets sequenced on BGISEQ-500, five datasets sequenced on DNBSEQ-G400) and two Illumina sequencing platforms (one dataset sequenced on HiSeq2500 and one dataset sequenced on NovaSeq6000), (Supplementary Table S2, See Methods). Each dataset was down-sampled and processed for BAM following our previous approach[20]. We got eight BAMs based on DNBSEQ™ platforms with average 31.43x and 99.02% whole genome coverage, and two BAMs based on Illumina platforms with average 30.60x and 99.19% whole genome coverage (Table 1). Then, these BAMs of ten datasets were used to detect CNVs across genome by five tools: BreakDancer, CNVnator, Pindel, DELLY and LUMPY, respectively. In total, we obtained 50 files of CNV results.

**Table 1.** Statistics of data and CNVs.

|  | Sequencing<br>strategy | Clean bases<br>(G) | Average<br>sequencing | Mapping<br>rate (%) | Duplicate<br>rate (%) | Coverage<br>(%) | BreakDancer | CNVnator | #CNV<br>Pindel | DELLY | LUMPY |
| --- | --- | --- | --- | --- | --- | --- | --- | --- | --- | --- | --- |
| BGISEQ-500_PE100 | PE100 | 113.6 | 37.44 | 99.22 | 2.45 | 99.12 | 2,329 | 5,778 | 4,924 | 1,904 | 2,378 |
| BGISEQ-500_PE150-1 | PE150 | 94.15 | 29.63 | 99.59 | 6.90 | 98.99 | 1,646 | 3,617 | 4,463 | 1,624 | 1,422 |
| BGISEQ-500_PE150-2 | PE150 | 94.20 | 29.51 | 99.57 | 7.49 | 98.97 | 2,265 | 4,241 | 4,310 | 2,209 | 2,202 |
| DNBSEQ-G400_PE100 | PE100 | 94.11 | 31.09 | 99.56 | 2.22 | 99.09 | 2,160 | 4,354 | 4,688 | 1,745 | 1,889 |
| DNBSEQ-G400_PE150_PCR-1 | PE150 | 94.11 | 30.83 | 99.54 | 2.19 | 98.99 | 1,576 | 2,234 | 4,577 | 1,902 | 1,609 |
| DNBSEQ-G400_PE150_PCR-2 | PE150 | 94.11 | 30.77 | 99.55 | 2.24 | 98.99 | 1,600 | 2,209 | 4,689 | 1,900 | 1,533 |
| DNBSEQ-G400_PE150_PCRfree-1 | PE150 | 94.11 | 31.13 | 99.60 | 1.24 | 98.98 | 1,519 | 2,890 | 4,221 | 1,705 | 1,463 |
| DNBSEQ-G400_PE150_PCRfree-2 | PE150 | 94.11 | 31.07 | 99.60 | 1.25 | 98.99 | 1,523 | 2,860 | 4,172 | 1,713 | 1,505 |
| HiSeq2500_PE150 | PE150 | 96.28 | 31.91 | 99.31 | 1.46 | 99.00 | 968 | 2,660 | 4,045 | 1,709 | 1,380 |
| NovaSeq6000_PE150 | PE150 | 93.69 | 29.30 | 99.06 | 5.91 | 99.06 | 897 | 1,229 | 5,730 | 1,773 | 1,350 |

Note: Pair-end (PE).

To reduce the false positives, firstly, low-quality CNVs were filtered out from raw CNVs (See Materials and Methods). Still, high ratio ambiguous CNVs were remained and these ambiguous CNVs might decreased the performance of CNV detection. Of which, a CNV in each CNV result was defined as ambiguous if it had at least 1bp overlap with other CNVs in the same CNV result, and the rest of CNVs were defined as unambiguous CNVs. We found average 9.82% (rang from 3.75% to 20.36%) CNVs detected by BreakDancer for ten datasets were ambiguous, while the ratio of ambiguous CNVs was 39.43% (range from 30.97% to 49.96%) for Pindel, 19.87% (range from 16.55% to 26.81%) for DELLY, 30.23% (range from 23.73% to 43.67%) for LUMPY and 0% for CNVnator, respectively (Supplementary Figure S2a). Further, we calculated the precision of ambiguous CNVs and unambiguous CNVs with two benchmarks, respectively (See Materials and Methods). For LUMPY_2014 benchmark, the precision of unambiguous CNVs was significantly higher than that of ambiguous CNVs (average 51.17% of unambiguous CNVs vs. average 12.83% of ambiguous CNVs, p = 5.64e-13 with t-test, Supplementary Figure S2b). Similarly, for 1KG_2015 benchmark, we also found the precision of unambiguous CNVs was significantly higher than ambiguous CNVs (average 39.23% vs. average 9.08%, p = 1.11e-16 with t-test, Supplementary Figure S2c). Those results indicated the negative effect of ambiguous CNVs and suggested the filtering of ambiguous CNVs to improve performance of CNV detection.

In a word, after removing low-quality and ambiguous CNVs from raw CNVs, we got final average 2,586 CNVs (range from 897 to 5,778) on genome-wide using five tools for ten datasets (Figure 1). The data size, major alignment statistics and total CNV calls using five tools were summarized in Table 1 (more statistics of CNVs were list on Supplementary Table S3).

**Figure 1.**
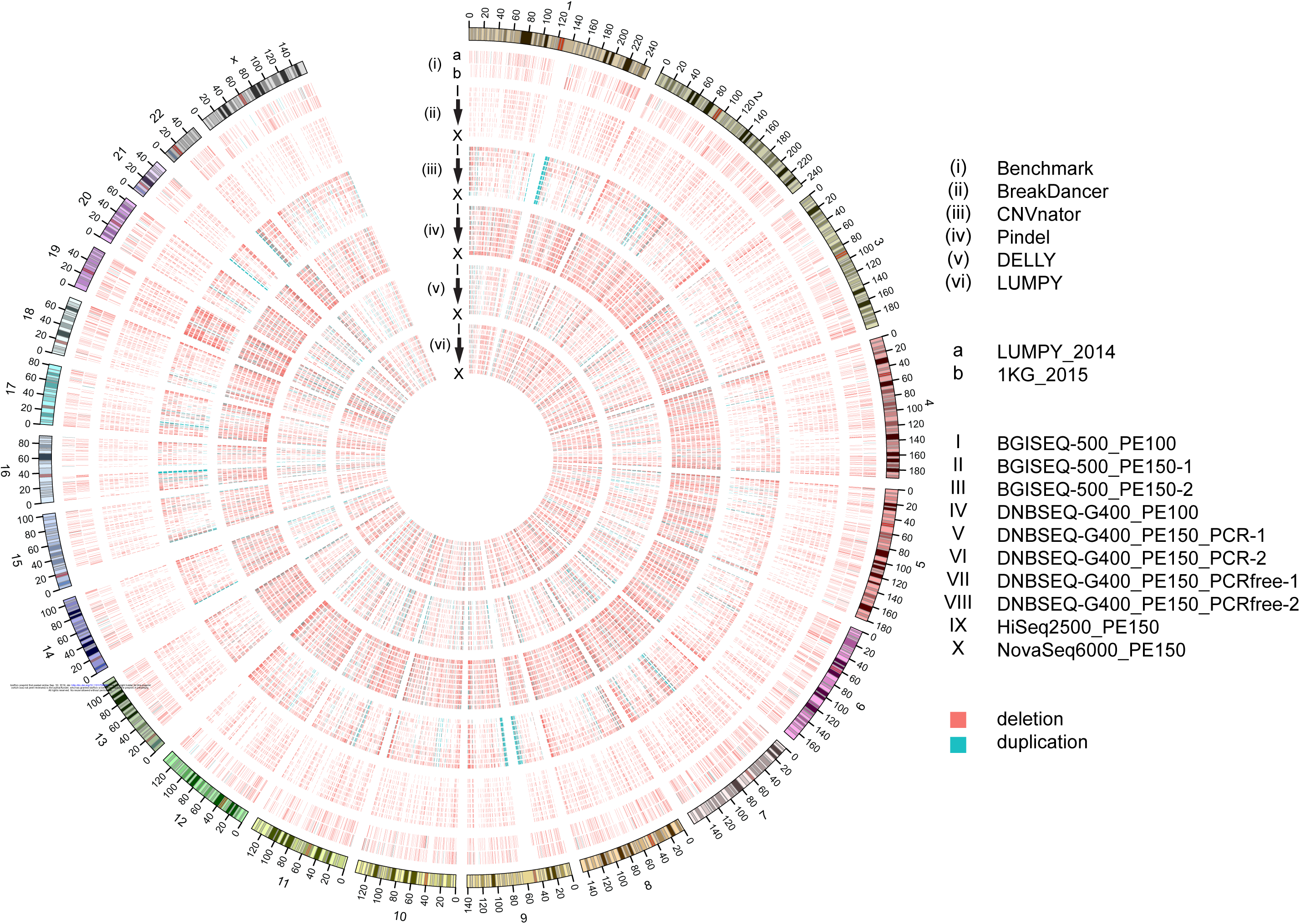
CIRCOS of CNVs. Circos plot depicting the landscape of CNVs on whole genome. The outermost circle shows the ideogram of chromosomes. The (i) circle displays the location of two benchmarks: LUMPY_2014 and 1KG_2015. The (ii), (iii), (iv), (v) and (vi) circle denote the distribution of CNV of ten datasets using BreakDancer, CNVnator, Pindel, DELLY and LUMPY, respectively.

### High quality CNVs produced from DNBSEQ™ platforms

To evaluate the performance of CNVs based on DNBSEQ™ platforms, two Illumina platforms were used as control platforms. Firstly, we compared the number of CNVs detected by each tool between DNBSEQ™ and Illumina sequencing datasets. We got average 1,827 (range from 1,519 to 2,329) CNVs using BreakDancer for eight datasets from two DNBSEQ™ platforms, while the CNV count was 3,523 (range from 2,209 to 5,778) using CNVnator, 4,506 (range from 4,172 to 4,924) using Pindel, 1,838 (range from 1,624 to 2,209) using DELLY and 1,750 (range from 1,422 to 2,378) using LUMPY (Table 1). While in two datasets from two Illumina platforms, we got 933 (897 and 968) CNVs using BreakDancer, 1,945 (1,229 and 2,660) using CNVnator, 4,888 (4,045 and 5,730) using Pindel, 1,741 (1,709 and 1,773) using DELLY and 1,365 (1,350 and 1,380) using LUMPY (Table 1). We found the average CNVs from DNBSEQ™ platforms was 1.96-fold of average CNVs from Illumina platforms using BreakDancer, 1.81-fold using CNVnator, 0.92-fold using Pindel, 1.06-fold using DELLY and 1.28-fold using LUMPY (Table 1). Those results showed that WGS data sequenced on DNBSEQ™ platforms could produce more CNVs on genome wide.

Secondly, we annotated CNVs by locating CNVs to seven genomic regions: Upstream, Exonic, Intronic, Downstream, Intergenic, CpG island (CGI) and CGI-shore. We found similar distribution of all 50 CNV result files (average 37.28% CNVs based on DNBSEQ™ platforms and 36.25% CNVs based on Illumina platforms were in Intergenic, 28.32% and 31.47% in Exonic, 27.49% and 25.19% in Intronic, 3.53% and 3.56% in Downstream, 3.38% and 3.53% in Upstream, 5.01% and 4.53% in CGI-shore,2.85% and 2.05% in CGI, Supplementary Figure S6, Supplementary Table S5), which suggested CNVs based on DNBSEQ™ platforms were highly consistent with Illumina platforms.

Further, we evaluated the sensitivity of CNVs based on DNBSEQ™ platforms and Illumina platforms. When compared with LUMPY_2014 benchmark, the average sensitivity was 54.44% of CNVs based on DNBSEQ™ platforms vs. 26.61% of CNVs based on Illumina platforms using BreakDancer, while 13.36% vs. 12.73% using CNVnator, 67.06% vs. 47.62% using Pindel, 38.64% vs. 30.08% using DELLY and 43.52% vs. 29.57% using LUMPY when compared with LUMPY_2014 (Figure 2, Supplementary Figure S5). And when the benchmark 1KG_2015 was used, the average sensitivity was 51.57% vs. 30.26% using BreakDancer, 25.98% vs. 24.53% using CNVnator, 70.47% vs. 52.85% using Pindel, 39.16% vs. 33.21% using DELLY and 42.96% vs. 32.66% using LUMPY (Figure 2, Supplementary Figure S5). Those results suggested that there were higher sensitivities of CNVs based on DNBSEQ™ platforms than CNVs based on Illumina platforms using different tool and different benchmark.

**Figure 2.**
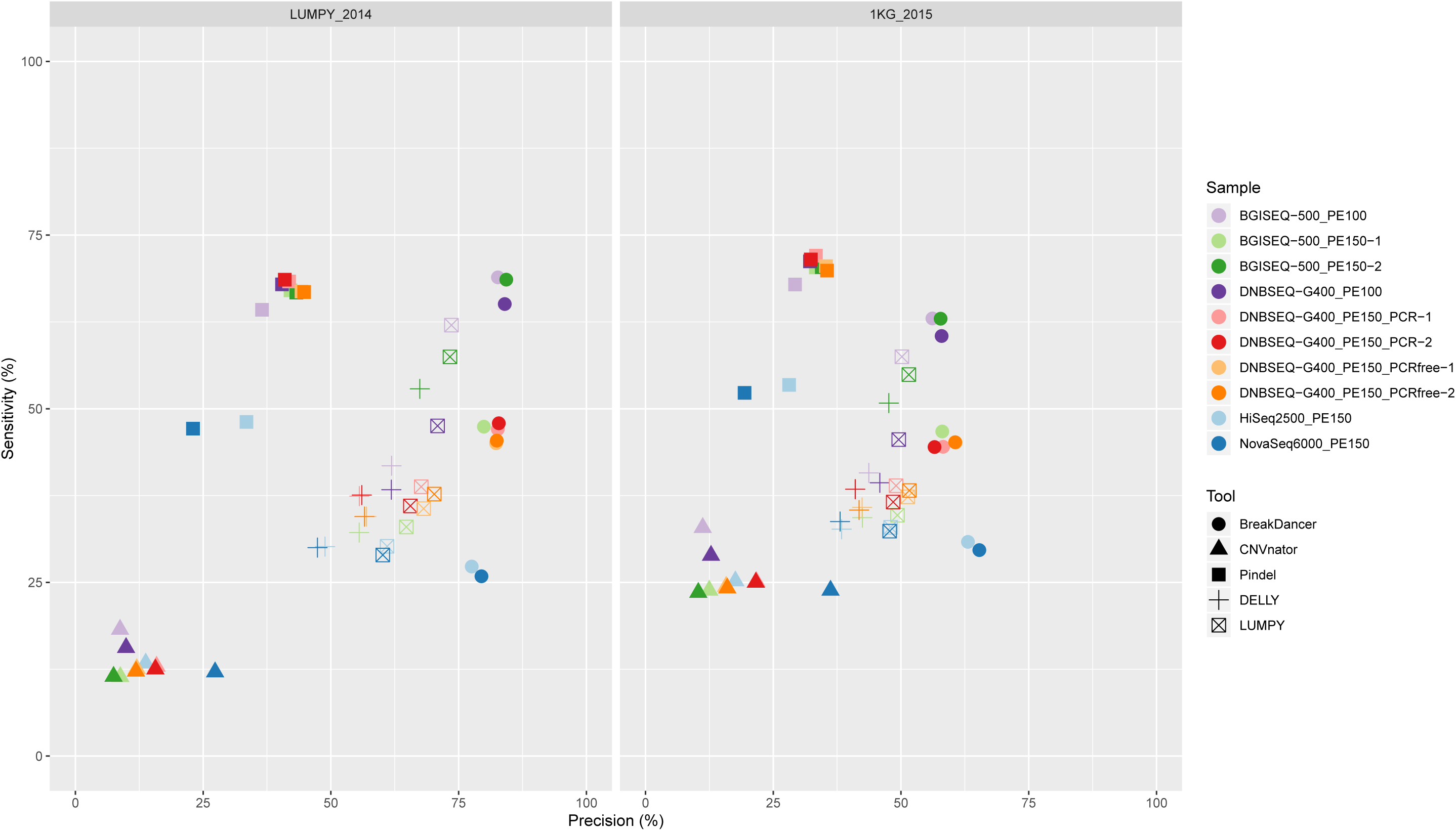
Evaluation of CNVs. Dot plot shows the sensitivity and precision of ten datasets using five tools by compared with LUMPY_2014 (left) and 1KG_2015 (right). Ten datasets are marked with different colors, and five tools are marked with different point types.

Meanwhile, we evaluated the precision of CNVs between DNBSEQ™ platforms and Illumina platforms and found the average precision was 82.67% vs. 78.53% using BreakDancer, 11.29% vs. 20.55% using CNVnator, 41.77% vs. 28.25% using Pindel, 58.97% vs. 48.12% using DELLY and 69.27% vs. 60.58% using LUMPY when compared with LUMPY_2014. While 58.26% vs. 64.22% using BreakDancer, 15.24% vs. 26.90% using CNVnator, 33.23% vs. 23.75% using Pindel, 43.23% vs. 38.23% using DELLY and 50.13% vs. 47.88% using LUMPY comparing with 1KG_2015. Those results showed that with two benchmarks, there were higher precisions of DNBSEQ™ platforms using Pindel, DELLY, LUMPY, but lower precisions of DNBSEQ™ platforms using CNVnator.

### Tool affects CNV detection

To investigate the characteristic of CNV detection by different tools, we firstly compared the number or the length of CNVs detected by different tools from the same datasets. We found that there were average 4,582 (range from 4,045 to 5,730) CNVs from ten datasets using Pindel, which 87.01% (39,869 / 45,819) CNVs were smaller than 1,000bp (Table 1, Supplementary Figure S3), verified that Pindel is an effective tool for small CNVs[15]. Meanwhile, we got average 3,207 (range from 1,229 to5,778) CNVs using CNVnator, which 95.78% (30,717 / 32,072) were larger than 1,000bp (Table 1, Supplementary Figure S3), showed that CNVnator with RD-strategy is more suitable to detect large CNVs. About BreakDancer, DELLY and LUMPY, we got CNVs with similar average CNV count (average 1,648, 1,818 and 1,673 CNVs for BreakDancer, DELLY and LUMPY, respectively) and similar size distributions which were major range from 100 to 5,000bp (Table 1, Supplementary Figure S3).

Secondly, we compared the sensitivity and precision between different tools. As the sensitivity and precision of CNVs detection described above, we found BreakDancer showed the highest precision but unstable sensitivity, while CNVnator had both lowest precision and sensitivity (Figure 2). Meanwhile, Pindel showed higher sensitivity but lower precision, while DELLY and LUMPY had both balanced sensitivity and precision (Figure 2).

Further, we studied the region distribution of CNVs across genomic from different datasets by different tools. We found CNVnator could detected more CNVs in Exonic, CpG Island (CGI) and CGI-shore than other tools (average 37.04% with 1,150 CNVs in Exonic, 5.43% with 198 in CGI and 8.84% with 291 in CGI-shore, Supplementary Figure S6, Supplementary Table S5), while BreakDancer and Pindel could detected more CNVs in Intronic and Intergenic (average 29.36% and 38.58% of CNVs in Intronic and Intergenic using BreakDancer, 32.03% and 39.00% using Pindel). No visible bias appeared in CNVs detected by DELLY or LUMPY.

In a word, we found similar results CNV count and length when analyzing different datasets with the same tool but variety when analyzing the same dataset with different tools. And distributions of length and genomics regions and performance would be helpful for more critical selection of CNV detection tool.

## Discussion

Detecting CNV using WGS sequenced on Illumina platforms were widely used and many tools were designed based on those requirements, but the feasibility of CNV detection on DNBSEQ™ platforms is unclear. This study represents a first systematic investigation and characterization of CNV detection based on DNBSEQ™ platforms and we found that both the sensitivity and the precision of CNVs based on DNBSEQ™ platforms were consistent with those of CNVs based on Illumina platforms, if not better. Furthermore, by comparing different CNV tools, we found that there were different features of CNV detected by different tools, such as CNV count, the length region distribution across genome, CpG Island (CGI), CGI-shore, sensitivity and precisionof CNVs, indicating that this study may help researchers to choose tools according to different conditions.

There are some critical parameters affect the performance of CNV detection. First parameter is the choice of CNV detection tools. Dozens of CNV detection tools were available and designed for different sample conditions. We chose five common tools for CNV detection on single WGS and still got obvious differences. BreakDancer built on RP could only detect mediocre deletions influenced by the insert size of library. CNVnator built on RD gets large CNVs with low sensitivity and precision. Pindel build on SR would get high proportion overlapped CNVs with worse precision. Those results indicated deficiency of current CNV detection tools and complicated tool selection for CNV detection.

The second parameter is the lack of a high-quality benchmark. We used two public benchmarks of NA12878 here. However, high proportion specific CNVs between benchmarks and opposite comparisons with two benchmarks between sensitivity and precision showed the influence of benchmark for CNV detection, and the necessity of a high-quality benchmark.

Finally, the uncertain saturation is another parameter in evaluation performance of CNV detection. A combination of current tools, benchmarks and data size could confuse the saturation and performance of CNV detection.

In summary, the various performance of CNV detection on DNBSEQ™ platforms indicate the situation of CNV detection and the need for developing more efficient and precise CNV detection and evaluation methods. CNV detection tools with high performance and efficiency can enhance CNV researches using WGS sequenced on NGS.

## Conclusions

WGS sequenced on NGS is an efficient approach for genome researches. The applies of WGS sequenced on Illumina platforms were widely-used for CNV detection with numerous achievements. However, the feasibility of CNV detection on DNBSEQ™ platforms were not clear. Our results indicate CNV detection using WGS sequenced on DNBSEQ™ platforms is accurate and efficient. Our study provides a comprehensive guide for CNV researchers using DNBSEQ™ platforms with tool selection, benchmark, performance.

## Materials and Methods

### Public data used

All fastq data of NA12878 were downloaded from website: Gigadb, NCBI and CNSA (Supplementary Table S2). All data were down-sampled for approximately 30x and processed following out previous WGS approach.

Two public benchmarks of CNVs of NA12878: 2,819 CNVs from Layer, et al. in 2014 (LUMPY_2014)[27] and 2,171 CNVs from the 1000 Genome Project WGS data in 2015[28] were used. Both benchmarks were extracted from EXCEL and processed with BEDtools (v2.16.2).

### CNV detection using BreakDancer

BreakDancer (ver.1.4.5) was used for CNV calling with default parameters with the read-pair strategy. Raw CNVs from BreakDancer were filtered out according to: (1) SV type is not ‘DEL’, (2) confidence score < 90, (3) supporting read pairs < 3, (4) not autosomal or chrX, (5) overlap a gap in the reference genome,

### CNV detection using CNVnator

CNVnator (ver.0.3.3) was used for CNV detection using a read-depth strategy. The optimal bin size for each data was chosen according to the authors’ recommendations, such that the ratio of the average read-depth signal to its standard deviation was between 4 and 5 (Supplementary Table S4). Raw CNVs from CNVnator would be filtered by those factors: (1) q0 ≥ 0.5 or q0 < 0, (2) e-val1 ≥ 0.05, (3) not autosomal or chrX, (4) overlap a gap in the reference genome,

### CNV detection using Pindel

Pindel (ver.0.2.5b9) was used with default parameters with the split read strategy. Raw CNVs from Pindel were filtered with SVTYPE is not ‘DEL’ or ‘DUP:TANDEM’ and supporting read pairs < 3. Also, CNVs with: (1) not autosomal or chrX, (2) overlap a gap in the reference genome were filtered out.

### CNV detection using DELLY and LUMPY

Both DELLY (ver.0.7.8) and LUMPY (ver.0.2.13) were used to detect CNVs with default parameters with a combination of approaches. CNVs from DELLY were filtered out if: (1) SVTYPE is not ‘DEL’ or ‘DUP’, (2) FILTER is ‘LowQual’, (3) supporting read pairs < 3. CNVs from LUMPY were filtered out if: (1) SVTYPE is not ‘DEL’ or ‘DUP’, (2) PE < 3. Also, CNVs with: (1) not autosomal or chrX, (2) overlap a gap in the reference genome were filtered out.

### Filtration of raw CNVs

Firstly, a raw CNV detected by tool was filtered for removing low-quality CNV. Then, a CNV was defined as ambiguous or unambiguous if it had at least 1bp overlap with other CNVs in the same CNV result or not. All ambiguous CNVs were filtered out for decreasing false positive.

### Evaluation of CNVs

The evaluation of CNV compared with benchmark was processed by in-house scripts. A CNV was considered valid if either it overlapped a single CNV in benchmark by ≥ 50% reciprocally in size, or there existed a set of CNVs in benchmark such that each CNV in benchmark had ≥ 50% size overlapped with the CNV and ≥ 50% of the CNV overlapped with this set of CNVs in benchmark. The sensitivity or precision were calculated using the following equations:

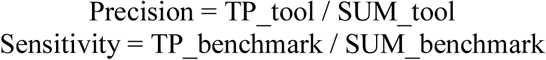

where TP_ tool was the number of validated CNVs, SUM_tool was the total number of CNVs, TP_ benchmark was the number of a set of CNVs in benchmark that overlapped with validated CNVs, and SUM_ benchmark was the number of CNVs in benchmark.

### Annotation of CNVs

Genomic regions and CGI were downloaded from UCSC (http://hgdownload.soe.ucsc.edu/goldenPath/hg19/database/). The region of 2,000bp frank of CGI in each direction were defined as CGI-shore. CNVs were located to any of seven regions (Upstream, Exonic, Intronic, Intergenic, Downstream, CGI and CGI-shore) if the midpoint of CNV region was in any regions.

## Authors’ contributions

J.R., L.P., X.L. and F.M. designed the project. J.R. and L.P. carried out the data analysis and wrote the manuscript, with critical comments and suggestions from the rest of the authors. All authors read and approved the final manuscript.

## Acknowledgements

This research was supported by the National Key R&D Program of China (2017YFC0906501), the Key R&D Program of Guangdong Province (2019B020226001).

## Additional files

Supplementary Figure S1. Venn of benchmarks. Venn showing the number and ratio of specific CNVs in each benchmark with 90.00% threshold (a) or 50.00% threshold (b).

Supplementary Figure S2. Summary of the distribution and precision of ambiguous CNVs. (a) shows the proportion distribution of ambiguous in 50 CNV results of ten datasets using five tools after low-quality CNV filtration. The comparison of precision between ambiguous and unambiguous CNVs with LUMPY_2014 (b) and 1KG_2015 (c) is display bellow.

Supplementary Figure S3. Length distribution. The distribution of CNV length for ten datasets using five tools.

Supplementary Figure S4. Evaluation between benchmarks. Comparison of sensitivity (a) and precision (b) between two benchmarks.

Supplementary Figure S5. Evaluation between platforms. Comparison of sensitivity and precision between Illumina platforms and DNBSEQ™ platforms with two benchmarks.

Supplementary Figure S6. Annotation of CNVs. Histogram shows the location of CVNs in genomic regions, CGI and CGI-shore. CpG Island: CGI. CpG island-shore: CGI-shore.

Supplementary Table S1. Tools for CNV detection.

Supplementary Table S2. Demo data.

Supplementary Table S3. Statistics of CNVs.

Supplementary Table S4. Bin size in CNVnator.

Supplementary Table S5. Annotation of CNVs.

