## Supplementary figures and images for "Detection and characterization of copy number variants based on whole-genome sequencing by DNBSEQ platforms"

### Supplementary Figure S1

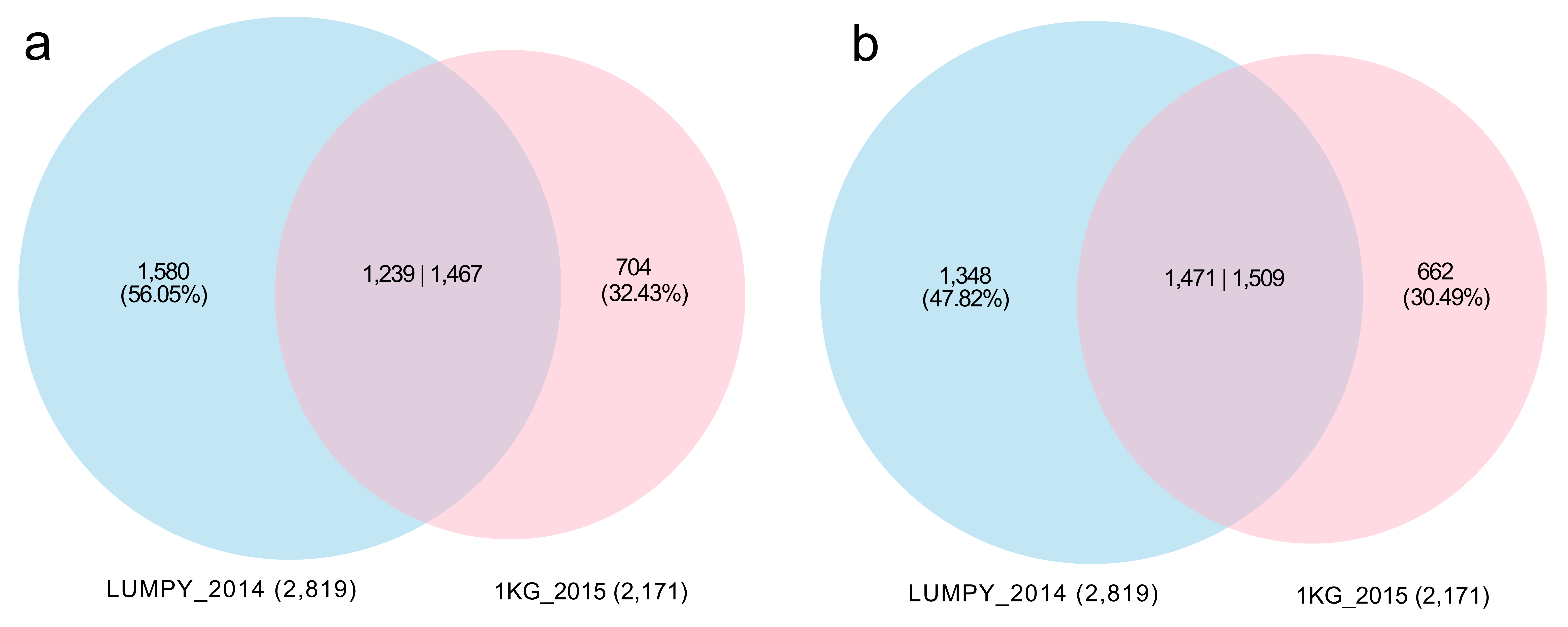

### Supplementary Figure S2

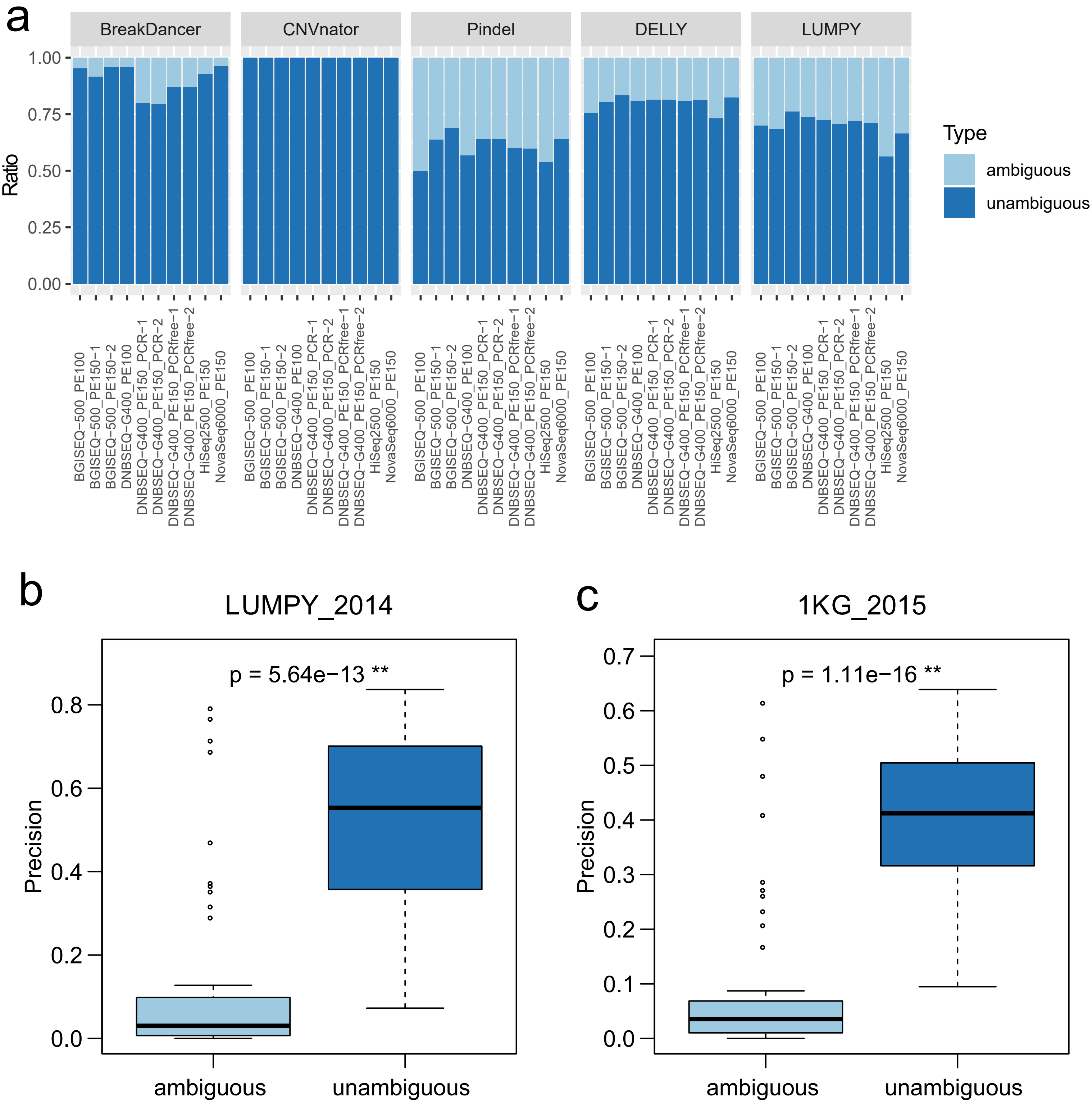

### Supplementary Figure S3

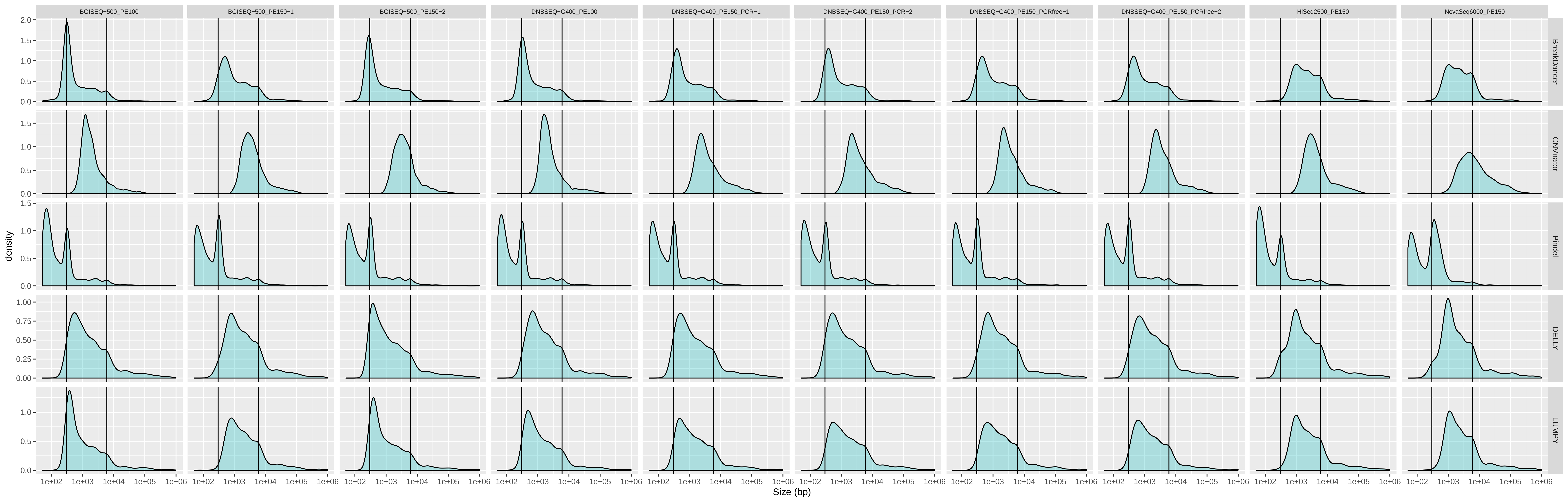

### Supplementary Figure S4

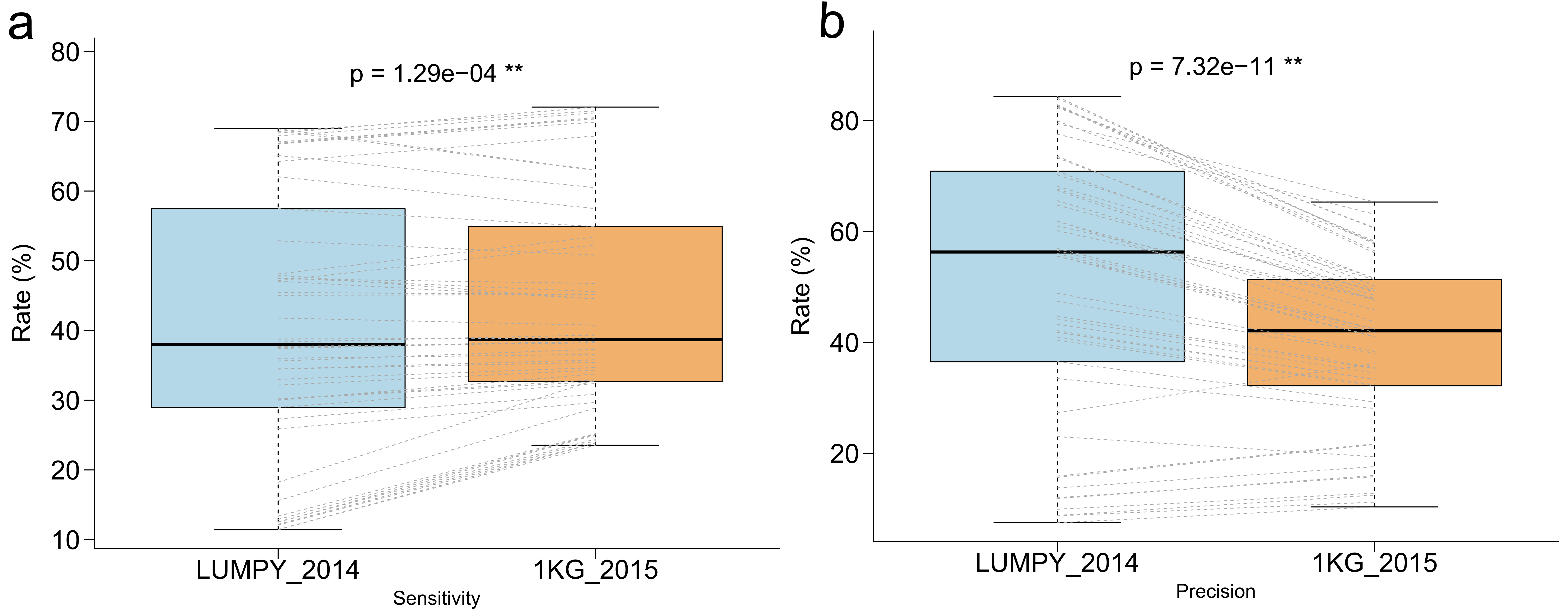

### Supplementary Figure S5

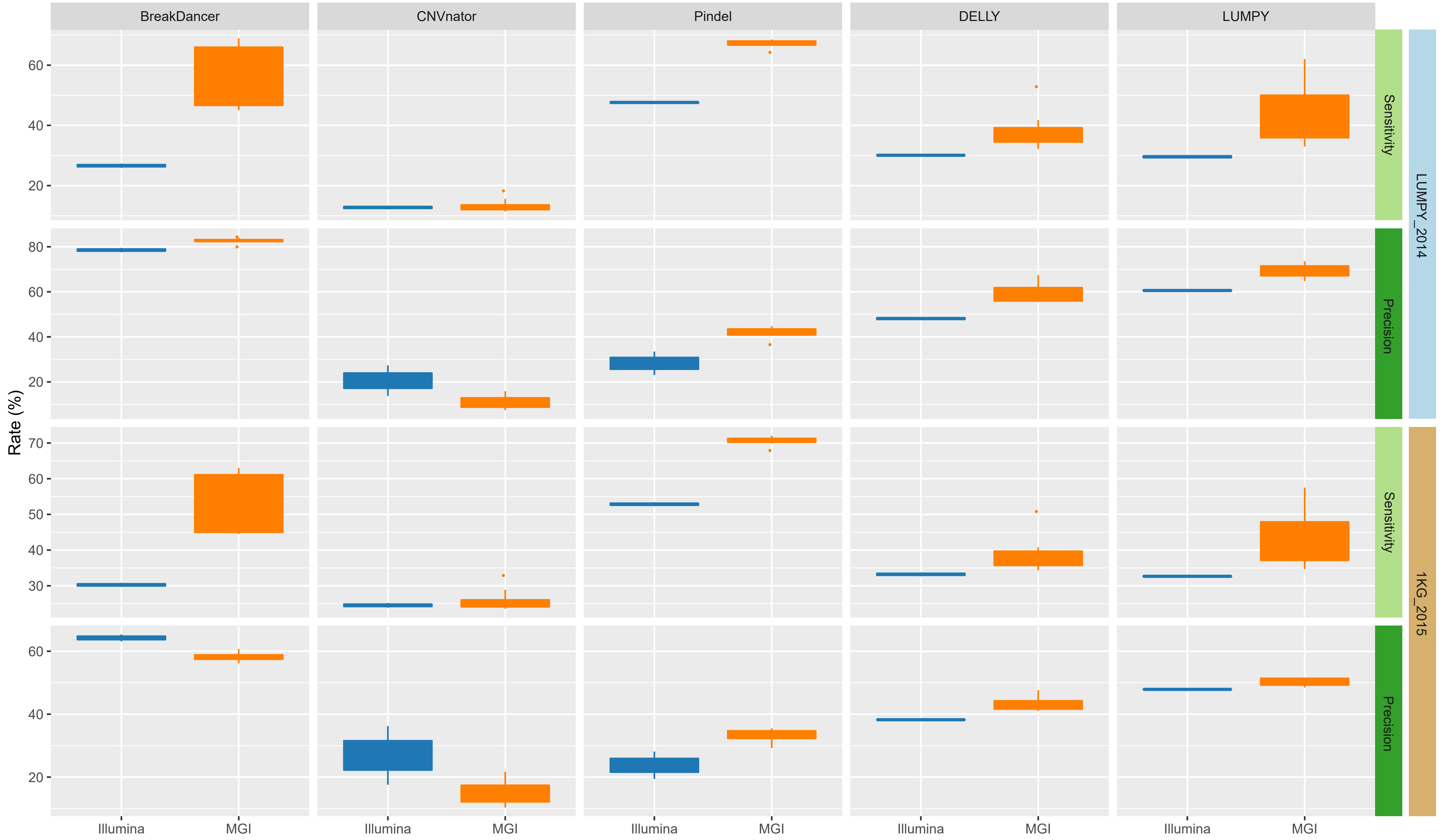

### Supplementary Figure S6

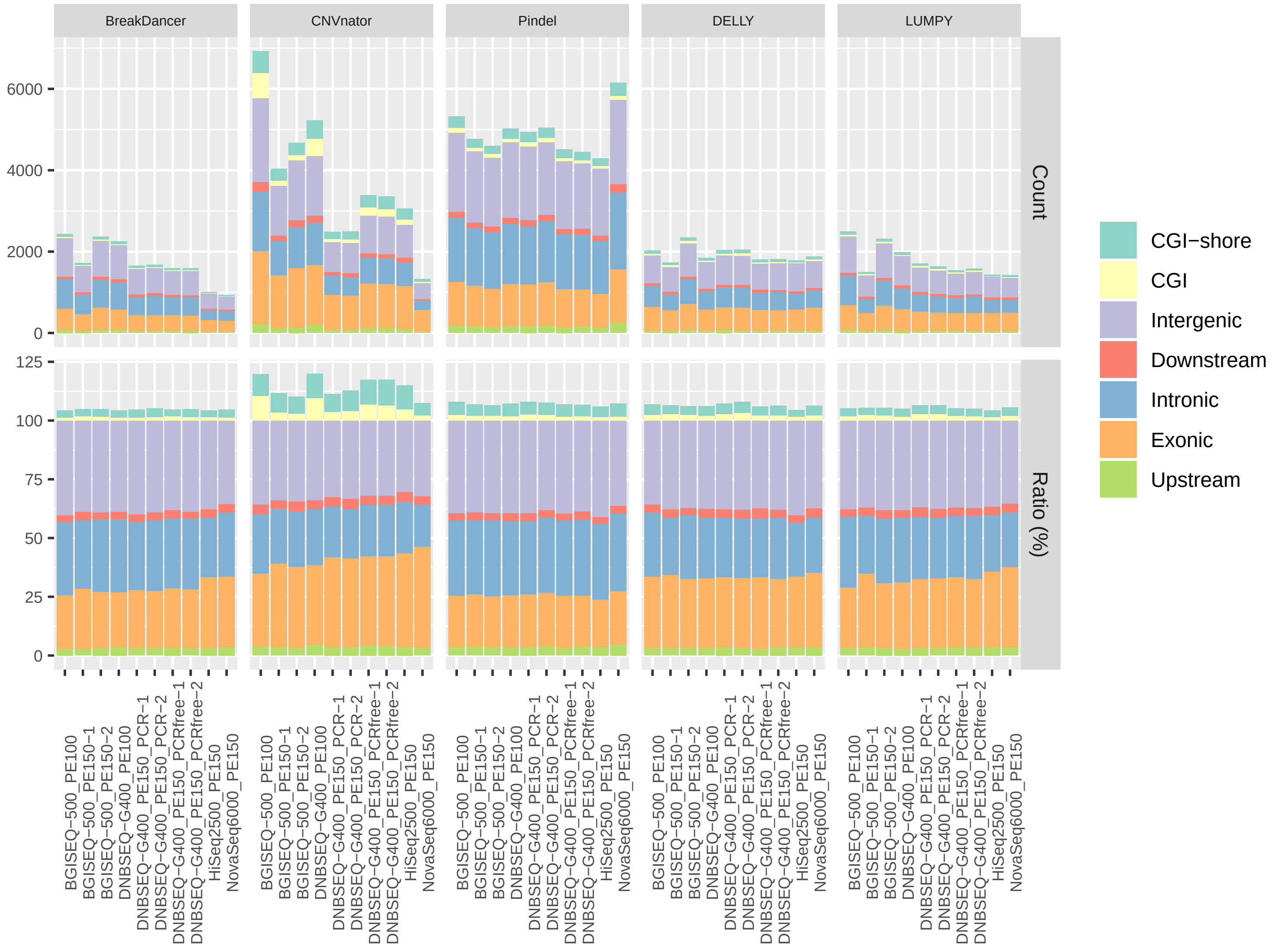
